# Oxygen/glucose-deprivation causes long-term impairment of synaptic CaMKII movement

**DOI:** 10.1101/2025.03.01.640973

**Authors:** Olivia R. Buonarati, Nidia Quillinan, K. Ulrich Bayer

## Abstract

Learning and memory are thought to require hippocampal long-term potentiation (LTP), a form of synaptic plasticity that is persistently impaired after cerebral ischemia and that requires movement of the Ca^2+^/calmodulin-dependent protein kinase II (CaMKII) to excitatory synapses. We show here that oxygen/glucose-deprivation (OGD) in cultures hippocampal neurons causes a long-lasting impairment of CaMKII movement. Notably, CaMKII inhibition at 30 min after onset of OGD prevented the impairment in CaMKII movement. Thus, CaMKII mediates both, LTP mechanisms and their ischemia-induced impairment. These findings provide a mechanism by which ischemic conditions can impair LTP and explain how CaMKII inhibition after cerebral ischemia can prevent these LTP impairments.

## INTRODUCTION

Cerebral ischemia, i.e. the lack of oxygen in the brain, causes lasting damage in several distinct but related conditions, including focal cerebral ischemia (stroke) and global cerebral ischemia (GCI)^1-4^. The most common form of stroke is caused by locally formed blood clots and is termed ischemic stroke; embolic stroke is caused by the lodged blot clots that were initially formed elsewhere, and hemorrhagic stroke is caused by blood vessel rupture^5^. GCI can be caused by drowning, suffocation or, most commonly, by cardiac arrest^6^. The immediate damage of any type of cerebral ischemia is acute neuronal cell death. In stroke, the area of cell death depends largely on the specific focal location to which blood flow is impaired. In GCI, the neurons most sensitive to cell death appear to be pyramidal neurons of the hippocampus^7-9^, a brain structure centrally involved in learning and memory^10,11^. However, in addition to causing cell death in a subset of neurons, the surviving hippocampal neurons show long-lasting functional impairments, specifically in long-term potentiation (LTP)^12-14^, a form of synaptic plasticity thought to be required for learning and memory^15-17^. Indeed, most GCI patients suffer from impairments in learning and memory^18-21^. It is likely that these learning and memory impairments are caused by a combination of the neuronal cell death and the long-lasting functional impairments in GCI patients. Notably, mouse models of GCI show that the functional impairments of LTP is an independent event, and not just a secondary result of neurons dying: The LTP impairments are still seen at 7 and even 30 days after GCI, i.e. at time points when any dead or dying neurons are completely removed and thus can no longer contribute to the impairments^12-14^. Additionally, inhibition of TRPM2 channels restored LTP even when done at delayed time points after the initial neuronal cell death had already occured^13^. The mechanisms of ischemic neuronal cell death have been studied intensively for decades, but this has not translated to therapeutic success. By contrast, much less is known about the mechanisms underlying the long-term functional impairments in LTP after cerebral ischemia^6^. Here, we used the hippocampal cell culture model of oxygen glucose deprivation (OGD) in order to study mechanisms of ischemia-induced LTP impairments. Specifically, we investigated the effect of OGD on the movement of the Ca^2+^/calmodulin-dependent protein kinase II (CaMKII) to excitatory synapses in response to LTP-inducing stimuli. This CaMKII movement is mediated by binding to the NMDA-type glutamate receptor GluN2B^22^, which is required for normal LTP^23,24^. We found that this CaMKII movement was persistently impaired by transient OGD, even 7 days after the OGD. Notably, like LTP in hippocampal slices on day 7 after GCI, the movement of CaMKII in hippocampal neurons was rescued by CaMKII inhibition with tatCN19o after the OGD. These results provide a mechanistic explanation for the LTP impairments after ischemia as well as for the rescue of such impairments by CaMKII inhibition.

## RESULTS

### Testing the long-term effect of OGD on CaMKII movement

OGD was induced in cultured rat hippocampal neurons on day *in vitro* (DIV) 10 using an anaerobic incubator, as detailed in the methods section. An OGD time of 1 h resulted in massive neuronal cell death. However, when OGD time was reduced to 30 min, a substantial number of neurons survived, allowing analysis of CaMKII movement on day 7-9 after OGD, i.e. on DIV17-19. Movement of endogenous CaMKII was monitored using intrabodies, as we have described previously^25^. For this, neurons were co-transfected 2-3 days before imaging with constructs coding for a YFP2-labelled intrabody directed against CaMKII and two mCherry- and mTurquois-labelled intrabody against marker proteins of excitatory versus inhibitory synapses, PSD95 and gephyrin, respectively. Then, synaptic localization of CaMKII was measured as ratio of CaMKII co-localized with PSD95 or gephyrin over CaMKII localized in the dendritic shaft. Movement of CaMKII was induced with chemical LTP stimuli (cLTP; 100 µM glutamate and 10 µM glycine for 45 sec). As expected from LTP stimuli, such cLTP increases surface expression of AMPA-type glutamate receptors^23,26,27^ and induces movement of CaMKII to excitatory but not inhibitory synapses^22,25,28^. The timeline of this experimental paradigm is illustrated in Figure 1A.

**Figure 1:**
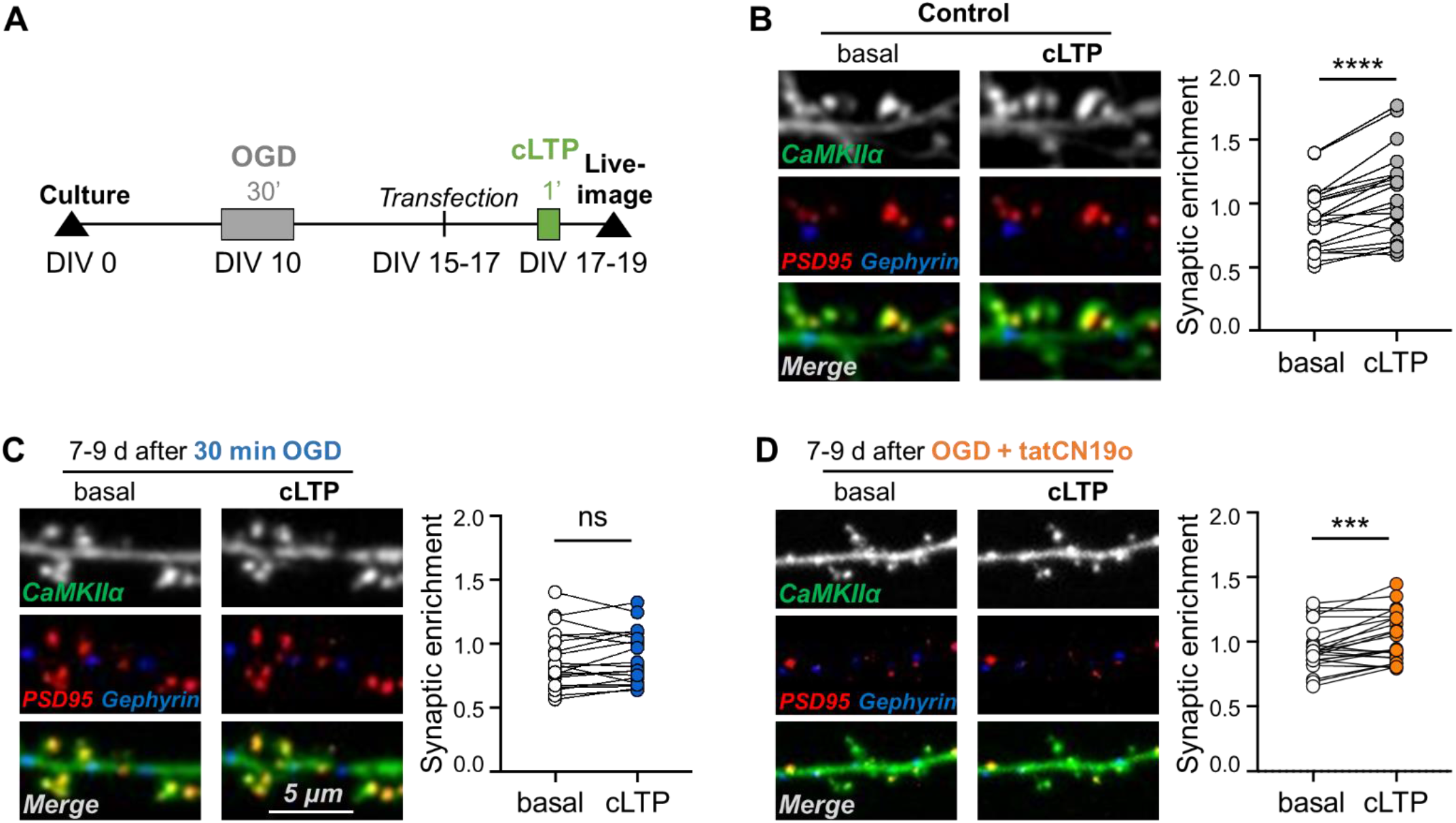
CaMKII movement is impaired by OGD and rescued by tatCN19o. (A)Timeline of CaMKII imaging after OGD in hippocampal neurons. On day *in vitro* (DIV) 10, rat hippocampal cultures were exposed to different durations of OGD. On DIV 15 to 17, neurons were transfected with constructs expressing intrabodies against CaMKII and maker proteins for excitatory versus inhibitors synapses. Two days later, on DIV 17 to 19, synaptic CaMKII localization was assessed before and after chemical LTP stimuli (cLTP; 100 μM glutamate, 10 μM glycine, 45-60 sec) to assess CaMKII movement to excitatory synapses. (B)As expected, cLTP stimulates movement of CaMKII to excitatory synapses (paired, two-tailed t-test, ****p<0.0001, n=21 cells). (C)However, 30 min OGD blocks CaMKII movement to excitatory synapses 7 to 9 days after the insult (paired, two-tailed t-test, ns p=0.3188, n=20 cells). (D)Application of CaMKII inhibitor tatCN19o (1 µM) restores CaMKII movement when applied 30 min after OGD onset (paired, two-tailed t-test, ***p=0.0006, n=24 cells).

### Transient OGD causes persistent impairment in CaMKII movement

As described previously, we found that cLTP stimuli induced a robust movement of endogenous CaMKII to excitatory synapses in hippocampal neurons (Fig. 1B). However, 7-9 days after exposing the neurons to 30 min of OGD, cLTP-induced CaMKII movement was completely abolished (Fig. 1C). When neurons were exposed instead to a briefer 15 min OGD, the cLTP-induced CaMKII movement appeared slightly reduced but was still significant (Supplemental Fig. S1A) and not significantly different from control without OGD treatment (Supplemental Fig. S1B). Thus, while 15 min OGD may have a mild effect on LTP-induced CaMKII movement to excitatory synapses, 30 min OGD completely abolished this CaMKII movement.

### The OGD-induced impairment of CaMKII movement is rescued by CaMKII inhibition

The LTP impairments after GCI are reversed by CaMKII inhibition with tatCN19o when applied 30 min after GCI^14^. As LTP requires CaMKII movement to excitatory synapses, we decided to test if tatCN19o also reverses the impairments in this CaMKII movement that is caused by OGD in hippocampal cultures. Indeed, CaMKII inhibition by addition 5 µM tatCN19o immediately after the OGD treatment (i.e. 30 min after onset of OGD) reinstated the cTLP-induced CaMKII movement 7-9 days after OGD to the same level as seen in control conditions without (Fig. 1D). While this rescue of CaMKII movement by CaMKII inhibition may appear surprising, this was the expected results based on the rescue of LTP by similar CaMKII inhibition after GCI *in vivo*^14^.

## DISCUSSION

This study demonstrated that (i) OGD causes a long-lasting impairment in the LTP-related movement of CaMKII to excitatory synapses and that (ii) this impairment is alleviated by CaMKII inhibition with tatCN19o. As LTP requires the CaMKII binding to GluN2B that mediates the CaMKII movement to excitatory synapses, these findings of this study provide direct mechanistic explanations for (i) the long-lasting impairment of LTP after ischemia^12-14^ and (ii) the rescue of LTP by tatCN19o^14^ that have been described previously. The rescue of CaMKII movement by CaMKII inhibition also implies that acute CaMKII activation directly mediates the ischemia-induced impairment of its own movement at delayed timepoints. Even though this may seem surprising, there is precedent for such an effect: CaMKII also mediates a similar impairment of its own movement that is triggered by amyloid β (Aβ)^25,29^, one of the major pathological agents in Alzheimer’s disease that also impairs LTP in hippocampal slices^30-32^. Indeed, CaMKII inhibition prevents not only the Aβ-induced impairment of its own movement in hippocampal neurons, but also the Aβ-induced impairment of LTP in hippocampal slices^32^. Thus, even though CaMKII is most famous for its physiological role in LTP, it appears to also have a pathological role in LTP impairment related to both cerebral ischemia and Alzheimer’s disease. As we have recently reviewed^33^, this dual role of CaMKII in LTP and LTP impairment is a concept that is just emerging. However, dual roles of CaMKII in two opposing functions have been described previously, including in both neuronal cell death and survival^34^ as well as in two opposing forms of synaptic plasticity, LTP and long-term depression (LTD)^34^. The CaMKII mechanisms enabling the dual role in both LTP and LTD have well-characterized over the last decade^34-37^. By contrast, the mechanisms underlying the role of CaMKII also in LTP impairment still await to be elucidated. This study provides a first step in this direction and indicates that the LTP impairment related to Alzheimer’s disease and after ischemia may have at least some overlapping mechanisms.

## ACKNOWLEDGEMENTS

This work was supported by National Institutes of Health grants F32 AG066536 (to O.R.B.), R01 NS046072 (to N.Q.), R01 NS081248, R01 NS118786, and R01 AG067713 (to K.U.B.).

## AUTHOR CONTRIBUTIONS

O.R.B. performed experiments and analyses. N.Q. provided essential technical advice. O.R.B., N.Q. and K.U.B. conceived this study. K.U.B. wrote the first draft of the manuscript, with input from all authors on the final draft..

## INCLUSION AND DIVERSITY

We support inclusive, diverse, and equitable conduct of research.

**Supplemental Figure S1:**
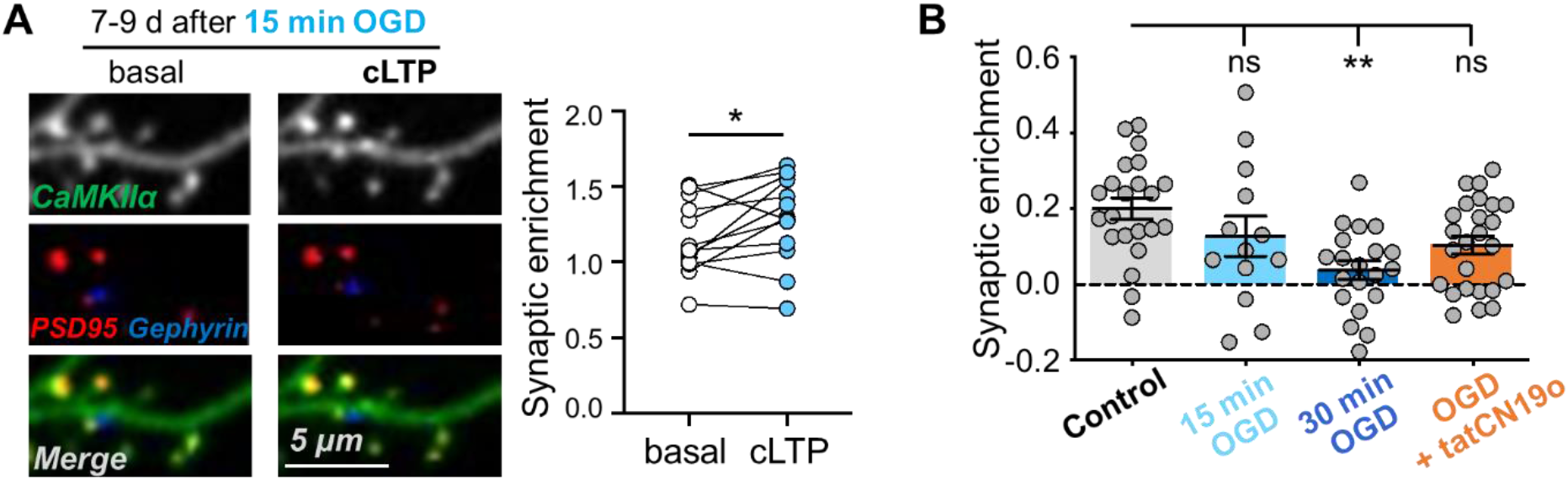
15 min of OGD has no significant effect on CaMKII movement. (A)In contrast to 30 min OGD (see Fig. 1B), 15 min OGD does not completely block cLTP-induced CaMKII movement to excitatory synapses on day 7-9 after the insult (paired, two-tailed t-test, *p=0.0322, n=13 cells). (B)Bar graph quantification shows the relative change in basal vs. cLTP after control (grey), 15 min OGD (light blue), 30 min OGD (dark blue), or 30 min OGD plus tatCN19o (orange) treatment. The change in synaptic enrichment of CaMKII is significantly lower after 30 min OGD, compared to control (one-way ANOVA with Tukey post-hoc test; control vs. 15 min OGD p=0.3422; control vs. 30 min OGD **p=0.0020; control vs. 30 min OGD + tatCN19o ns p=0.1217). Error bars indicate s.e.m.

## STAR METHODS

Key Resources Table

| REAGENT or RESOURCE | SOURCE | IDENTIFIER |
| --- | --- | --- |
| **Chemicals, Peptides, and Recombinant Proteins** |  |  |
| Papain | Worthington | LS 03126 |
| Lipofectamine 2000 | Invitrogen | 11668027 |
| B-27 supplement | Gibco | 17504044 |
| Penicillin-Streptomycin | Gibco | 15070063 |
| Neurobasal-A Medium | Gibco | 10888022 |
| MEM | Gibco | 11090081 |
| 5-Fluoro-2'-deoxyuridine (FdU) | Sigma | F0503 |
| Glutamate | Sigma | 6106-04-3 |
| Glycine | Sigma | 56-40-6 |
| tatCN19o | CPC Scientific;<br>Coultrap et al. 2011 | N/A (custom peptide) |
| **Recombinant DNA** |  |  |
| CaMKIIα-FingR-GFP | Dr. Donald Arnold (USC);<br>Mora et al, 2013 | N/A |
| PSD-95-FingR-GFP | Addgene | 46295 |
| Gephyrin-FingR-GFP | Addgene | 46296 |
| YFP-CaMKIIα | Addgene | 15214 |
| **Experimental Models: Cell Lines** |  |  |
| Primary hippocampal cultures | In House | N/A |
| **Experimental Models: Organisms/Strains** |  |  |
| Rat: Sprague-Dawley | Charles River Labs | N/A |
| **Deposited data** |  |  |
| Raw and analyzed data | This paper |  |
| **Software and Algorithms** |  |  |
| Slidebook 6.0 | Intelligent Innovations (3i) | RRID:SCR_014300 |
| Prism 7.0 | Graphpad | RRID: SCR_002798 |
| ImageJ | NIH | RRID:SCR_003070 |

## RESOURCE AVAILABILITY

### Lead contact

Further information and requests for resources and reagents should be directed to and will be fulfilled by the Lead Contact, K. Ulrich Bayer.

### Materials availability

Plasmids used in this work will be available upon request.

## EXPERIMENTAL MODEL AND SUBJECT DETAILS

All animal treatment was approved by the University of Colorado Institutional Animal Care and Use Committee, in accordance with NIH guidelines. Animals are housed at the Animal Resource Center at the University of Colorado Anschutz Medical Campus (Aurora, CO) and are regularly monitored with respect to general health, cage changes, and overcrowding. Pregnant Sprague-Dawley rats were supplied by Charles River Labs. Rat hippocampal cultures were prepared on postnatal day 0 (P0) and imaged on day *in vitro* (DIV) 17-19.

## METHOD DETAILS

### Material and DNA constructs

Material was obtained from Sigma, unless noted otherwise. The expression vectors for the GFP-labeled FingR intrabodies targeting CaMKIIα, PSD-95, and gephyrin were kindly provided by Dr. Donald Arnold (University of Southern California, Los Angeles, CA, USA) as described previously^38,39^. The FingRs against PSD95 and gephyrin contained the CCR5TC repressor ^38^; the FingR against CaMKIIα did not contain a repressor^39^. The fluorophore label was exchanged to contain the following tags in place of GFP: CaMKIIα-FingR-YFP2, PSD-95-FingR-mCh, and gephyrin-FingR-mTurquois^25^. These modifications were performed using Gibson Assembly^40^. The fluorescently-labelled intrabodies do not affect synaptic functions^25,38^. All constructs were validated by sequencing.

### Primary hippocampal culture preparation

To prepare primary hippocampal neurons, hippocampi were dissected from P0 mixed sex rat pups, dissociated in papain (7 mL HBSS buffered saline, 150 µL 100mM CaCl_2_, 10 µL 1M NaOH, 10 µL 500mM EDTA, 200 units papain) at room temperature for 1 h, followed by trituration. Cells were then plated at 100,000 cells/mL on poly-D-lysine (0.1 mg/mL in 1 M Borate Buffer) and laminin (0.01 mg/mL in PBS)-coated 18 mm glass coverslips in plating media (MEM, 10% FBS, 1% Penicillin-Streptomycin). On DIV1, plating media was replaced with feeding media (Neurobasal A containing 2% B27 and 1% Glutamax). On DIV7, half of the conditioned feeding media was replaced with fresh feeding media containing 2% 5-Fluoro-2’-deoxyuridine (FdU) to suppress glial growth.

### Oxygen Glucose Deprivation

OGD was induced in DIV10 neuronal cultures by replacing culture media with glucose-free deoxygenated saline solution containing (in mM): 140 NaCl; 5 KCl; 0.8 MgCl_2_; 1 CaCl_2_; 10 HEPES; 10 glucose, 10; pH 7.35 with NaOH) and placing the cultures in an anaerobic incubator containing 5% CO2/95% N2 at 37°C (Coy Laboratories Products, Grass Lake, MI, USA; contains catalyst ensuring oxygen levels <1 p.p.m.) for 15 or 30 min. Following OGD, cultures were returned to normoxia and normal culture media for subsequent transfection (DIV14-15) and imaging (DIV17-19). A subset of cultures were treated with 1 µM tatCN19o in the normoxia culture media immediately after (0 min) 30 min OGD.

### Image acquisition and analysis

DIV14-15 neuronal cultures were transfected with intrabodies at a 1:1:1 ratio, at a concentration of 1 µg total DNA/well, using Lipofectamine 2000. Two to three days after transfection, DIV 17-19 neurons were imaged using using an Axio Observer microscope (Carl Zeiss) fitted with a 63x Plan-Apo/1.4 numerical aperture (NA) objective, 50 μm pinhole, using 445, 515, 567 and 647 nm laser excitation and a CSU-XI spinning disk confocal scan head (Yokogawa) coupled to an Evolve 512 EM-CCD camera (Photometrics) and controlled using Slidebook 6.0 software (Intelligent Imaging Innovations [3i]. The 63x immersion objective (4.92 pixels/micron) was used to acquire all images.

Images were collected at 32°C in HEPES buffered imaging solution (ACSF) containing (in mM) 130 NaCl, 5 KCl, 10 HEPES pH 7.4, 20 Glucose, 2 CaCl_2_, 1 MgCl_2_. Hippocampal neurons were selected based on pyramidal shaped soma and presence of spiny apical dendrites, and tertiary dendritic branches were selected for analysis to maintain consistency. Images were acquired before and 5 min after stimulation. 2D maximum intensity projection images were then generated and analyzed using Slidebook 6.0. The average YFP intensity (CaMKIIα) at excitatory and inhibitory synapses was quantified using threshold masks of PSD-95 and gephyrin signal to calculate a puncta-to-shaft ratio (mean YFP intensity at PSD-95 or gephyrin puncta, divided by the mean YFP intensity of the adjacent dendritic shaft).

### Chemical LTP stimulation

In the imaging chamber, glutamate-induced chemical LTP (cLTP) was induced with 100 μM glutamate and 10 μM glycine for 45-60 s. Treatments were followed by washout with 5 volumes (5 mL) of fresh ACSF.

## QUANTIFICATION AND STATISTICAL ANALYSIS

All data are shown as mean ± SEM. Statistical significance is indicated in the figure legends. Statistics were performed using Prism (GraphPad) software. Imaging experiments were obtained and analyzed using SlideBook 6.0 software. All comparisons between two groups met parametric criteria, and independent samples were analyzed using unpaired, two-tailed Student’s t-tests. Comparisons between three or more groups meeting parametric criteria were done by one-way ANOVA with specific post-hoc analysis indicated in figure legends. Welch’s ANOVA was chosen due to its robustness against violations of the assumption of equal variances, ensuring reliable comparisons even when heterogeneity of variances exists across groups. Asterisks represent level of significance: *p<0.05; **p<0.01; ***p<0.001, ****p<0.0001.

